# New broad-spectrum anti-Flavivirus human antibodies with Zika virus-neutralizing potential

**DOI:** 10.1101/2022.04.01.486666

**Authors:** Renato Kaylan Alves de Oliveira França, Jacyelle Medeiros Silva, Lucas Silva Rodrigues, Andrea Queiroz Maranhão, Marcelo Macedo Brigido

## Abstract

Flavivirus infections show recurrent outbreaks and can be responsible for disease complications such as Hemorrhagic Dengue Fever and Congenital Zika Virus Syndrome. Effective therapeutic interventions are still a challenge. Antibodies can provide significant protection, although antibody response may fail due to ADE (Antibody-Dependent Enhancement) reactions or immune escape mutations. To generate effective neutralizing antibodies, the choice of the target antigen is a crucial part of the process. Human anti-Flavivirus antibodies were selected from a combinatorial library displayed on a phage surface. The antibodies were selected against a mimetic peptide based on the fusion loop region in Domain II of the protein E, which is highly conserved among different Flavivirus. Four rounds of selection were performed using the synthetic peptide in two strategies: the first was using acidic elution of bound phages, and the second was applying a competing procedure. After panning, the selected VH and VL domains were determined by combining NGS and bioinformatic approaches. Three different human monoclonal antibodies were expressed as scFvs and further characterized. All showed binding capacity to Zika (ZIKV), Yellow Fever (YFV), and Dengue (DENV) viruses. Two of these antibodies, AZ1p and AZ6m, could neutralize the ZIKV infection in a PRNT assay. These new antibodies have the potential to be used in therapeutic intervention against different Flavivirus illnesses and, due to the conservation of the fusion loop region, they may be resistant to scape mutations.

**Author summary:** The central idea of this work is to present a possible unique therapeutic approach to combat different diseases that cause health problems annually, such as Dengue and yellow fever infections and Zika congenital syndromes. The viruses that cause these diseases, of the Flavivirus genus, typically have disease amplification reactions and evasion mechanisms of the immune response that hinder the success of specific therapies and vaccines and require new forms of effective and safe treatments. We have developed new neutralizing human antibodies so that they bind to a highly conserved region of Flavivirus, called the fusion loop, and in such a way as to avoid adverse effects associated with anti-Flavivirus antibodies. We showed that the antibodies have high cross-reactivity against different Flaviviruses and we exemplified their neutralizing capacity for Zika virus infection in an ex vivo assay. New monoclonal antibodies such as those presented here may contribute to the control of important tropical diseases in a safer and more efficient way.

## Introduction

Flavivirus infections are a global health problem that cause relevant social and economic impacts in different countries, especially in the Americas, Africa, some European countries and Asia, causing death and illness of millions of people every year. Moreover, non-endemic areas are also in danger, with the potential for outbreaks due to climate change [1]. Nearly 1 billion people are at risk of contracting the dengue virus, and dengue cases have increased 30 fold in recent years [2]. This scenario is complicated by the resistance found in these viruses, such as the development of mutations that provide immunological escape or even by ADE (antibody-dependent enhancement) phenomenon that amplify the infection. Together, these factors are prejudicial to the success of therapies and vaccines [3].

Flavivirus disease outbreaks pose new challenges for clinics, mainly because of the onset of severe cases, with symptoms such as bleeding, shock and impairment of vital organs observed in hemorrhagic dengue fever [4]. Also, Zika virus infection is related to neurological impairments, such as Guillain-Barré syndrome, meningoencephalitis and congenital malformations, mainly due to the virus’ ability to infect progenitor neuronal cells [1,5,6]. The so-called Congenital Zika Virus Syndrome (CZS) has become a popular object of investigation. The Skeletal Muscular System, the Peripheral Nervous System of fetuses and the encephalon can be affected [7,8].

The *Flaviviridae* family comprises enveloped arboviruses whose genetic material is a positive-sense single-stranded RNA molecule. Different members of this family, such as Zika virus (ZIKV), Yellow Fever virus (YFV), Dengue virus (DENV), West Nile virus (WNV) and Japanese Encephalitis virus (JEV), cause critical human diseases. The viral RNA molecule encodes a polyprotein that is cleaved into three structural proteins: capsid protein (C), pre-membrane protein (prM) and envelope protein (E), and into seven non-structural proteins (NS1, NS2A, NS2B, NS3, NS4A, NS4B and NS5). These latter are involved in virus replication and assembly, and the inhibition of the antiviral immune response [9].

Protein E contains three domains: Domain I, which represents the N-terminal portion and influences viral tropism; Domain II, which comprises the dimerisation region and the fusion loop, and Domain III with a binding function to membrane receptors [10,11]. The fusion loop (FL) is the most conserved region among Flavivirus and plays a role in the infection process. After the virus binds to specific receptors, the viral particle is endocytosed. Due to the acidic pH in the late endosome, the viral envelope exposes the fusion loop, which binds to the endosomal membrane [12,13].

The immunotherapy of viral infections mediated by monoclonal antibodies is a relevant therapeutic approach. These molecules can block the viral infection cycle at different stages and induce an increase in the antigenic presentation and cellular immune responses [14,15]. Safe and efficient Flavivirus neutralizing antibodies are an alternative for vaccination for emerging viruses and immunocompromised people [16]. Human anti-Flavivirus antibodies are reported here. They were selected from a naive Phage Display Library by their ability to bind to a ZIKV fusion loop-derived peptide. The selected VH and VL domains were combined and expressed as scFv to characterize binding and neutralizing activities. The results pose these molecules as a potential treatment against different Flavivirus infections.

## Materials and Methods

### Cell lines and virus strains

*Escherichia coli* cells, lineage *XL1-BLUE MRF’* (Stratagene, cat: 200230), with resistance to Tetracycline, were grown in LB medium (Luria Bertani; 1.0% Peptone from casein, 0.5% Yeast Extract, 1.0% NaCl). This strain was used for the transformation, infection by phage and DNA extraction experiments. *Escherichia coli* cells, strain *Shuffle pLys Y* (New England Biolabs, cat: C3030J) compatible with pET vectors (DE3), were cultured in TB medium (Terrific Broth; 2.0% Peptone from casein, 2.4% Yeast Extract, 0.4% Glycerol, 72 mM K2HPO4, 17 mM KH2PO4). This strain was used for heterologous expression of recombinant antibodies.

Renal cells of green monkey *Cercopithecus aethiops, Vero* cells (ATCC, cat: CCL-81), were cultured in MEM medium (Eagle’s minimal essential medium; Thermo Fisher Scientific, cat: 61100061). Those cells were used for plaque reduction neutralization assays (PRNT) with the recombinant antibodies (described above). Zika virus PE243 strain (GenBank MF352141) was used for ELISA assays (Enzyme-Linked Immunosorbent Assay) and neutralization assays. Yellow Fever virus lineage 17DD (GenBank AF246798.1) and Dengue virus type 1-4 (Diagnostic kit anti-Dengue type 1-4 IgM, cat: EI 266a-9601-1 M, pre-coated plates, EUROIMMUN, Germany) were used for the ELISA assays.

### Antigen and antibody library preparation

The antigens were designed considering the protein E sequences of different Flavivirus, available in the GenBank database (DENV_1 BR/SJRP/1890/2018: MN631102.1; DENV_2 BR/SJRP/V2796/2019: MN631136.6; DENV_3 424/BR-PE/06: JX669508.1; DENV_4 Gu/SP/BR_1229: KP704217.1; WNV_H-442: AF459403.3; YFV_AR350397/Brazil/1979: U23570.1; JEV_JaOH0566: AY029207.1; ZIKV_LMM/AG5643: MT437401.1; ZIKV_PRI/PRVABC59_17/2015: MH916802.1; ZIKV_MR766: MW143022.1) and the PDB file 5IRE. Chimera was used to explore molecular structure [17]. A mimetic peptide to the Flavivirus fusion loop was designed, using the entire FL sequence, connected by a disulfide bridge to the SRCPT, a structurally associated region of domain II of the E protein of ZIKV. This peptide was synthesised conjugated to a biotin molecule (biotinylated peptide) to serve as a binding target for the fusion phages. A competitor peptide corresponding only to the fusion loop sequence was also designed. Synthetic peptides were manufactured by FastBio (Biomatik, Canada).

Electrocompetent *XL1-BLUE MRF’* cells (efficiency ∼1 × 10^9^ CFU/µg) were transformed, by electroporation, with a library of M13 filamentous phage protein III-conjugated human antibodies, with an estimated size of 1.7 × 10^8^, generated from B cells of human peripheral blood [18]. Electroporation was performed with 0.2 cm electrical cuvettes and with the following electrical parameters: 2.5 kV; 25 µF and 200 Ω. For library amplification, the transformed cells were cultured in SB medium (3.0% Bacteriological Peptone, 2.0% Yeast Extract, 1.0% MOPS, pH 7.0) containing 10^12^ pfu/mL helper phage VCSM13, 1.0% glucose, 10 µg/ml Tetracycline, 50 µg/ml Carbenicillin, 70 µg/ml Kanamycin, at 300 rpm and 37°C. On the following day, the fusion phages were precipitated by incubation of the 200 mL culture supernatant with 8 g of PEG-8000 and 6 g of NaCl, followed by 30 minutes incubation in ice bath centrifugation at 15,000g at 4 °C for 15 min. Phages were resuspended with 2 ml of 1.0% BSA solution (Bovine Serum Albumin, Sigma-Aldrich, cat: A7030) in TBS (50 mM Tris-HCl pH 7.5; 150 mM NaCl) and treated as input phage for selection.

### Library panning

Briefly, a 96-well flat-bottom NUNC PolySorp ELISA plate (Thermo Fisher Scientific, cat: 456529) was sensitised with 100 µL of 95 µM Streptavidin (Thermo Fisher Scientific, cat: 21122) in TBS for one hour at 37°C. The plate was washed three times with 200 µL of TBST (TBS with 0.1% Tween-20) and 100 µL of 1.7 µM viral antigen (biotinylated peptide) in TBS were added. The plate was incubated for one hour at 37°C and then washed. Blocking was performed with 150 µL of 3% BSA in TBST at 4 °C overnight. On the next day, 100 µL of input phages were added to the blocked plate, incubated for one and a half hours at 37°C.

Non-binding phages were removed with a variable number of washes at each selection round (5 and 10 washes for rounds 1and 2, respectively; 15 washes for rounds 3 and 4). The binding phages were eluted (output) through two elution strategies: one, using 100 µL of 100 mM Glycine-HCl solution pH 2.2, neutralized with 6 µL of neutralizing solution (Tris-base 2 M); and another, using a 100 µL solution of free-peptide (2.7 µM). The eluted phages were re-amplified for a new selection round, following the library amplification protocol described previously, in a culture final volume of 100 mL, transfecting an *E. coli* culture (2 mL) with 100 µL of phage eluate for 15 min at room temperature, instead of transformation.

### Analysis of the selection of specific antibodies

The heavy and light chain variable domains of rounds 0 and 4 (before and after both selections-acidic and competitive elution) were amplified with Platinum Taq DNA Polymerase High Fidelity kit (Thermo Fisher, cat:). The following oligonucleotide primers were used: sense oligonucleotide, LeadVH, (5’-CTGCCCAACCAGCCATGGCC-3’) and antisense oligonucleotide, VH_rev, (5’-CGATGGGCCCTTGGTGGAGGC-3’) for heavy chain, and sense oligonucleotide, Vkappa, (5’-GGGCCCAGGCGGCCGAGCTC-3’) and antisense oligonucleotide, VKappa_rev, (5’-AAGACAGATGGTGCAGCCACAGT-3’) for light chain. These oligonucleotides are specific for the regions of leader sequences (ompA and pelB) immediately before the V gene sequences and for the regions immediately after the J sequences of the variable domains [19].

The amplicons of VH and VL were sequenced using the Miseq system, with a 2×250 bp readout from the Illumina platform. The quality of the reads was verified with the FASTQC program, and the sequencing results were analysed with the automated immunoglobulin analysis tool, ATTILA, developed by the Molecular Immunology group at the University of Brasília [20]. In this tool, V(D)J signatures were determined, and the frequency variation of each sequence (fold-change) during selection was evaluated. Frequencies were calculated for each unique sequence with unique CDRs, and the fold-change (FC) value represents the changing the frequency of each sequence at the end of selection compared to its respective frequency before selection. ATTILA also assign V gene segment family based on the IgBLAST sequence bank, identifying the closest germline amino acid sequence.

The degree of dissimilarity was determined as the number of variations in the amino acid sequences of the most enriched antibodies. The length and the hydrophobicity of the amino acid CDR3 sequences of selected VH and VL were also analysed. Kyte and Doolittle hydrophobicity scale was used to calculate the GRAVY hydrophobicity scores (ProtParam Tool, Expasy; https://web.expasy.org/protparam/) [21].

### Recombinant antibody design, cloning and expression

The most enriched VH and VL in the selection were combined to construct single-chain antibody fragments (scFv). These recombinant antibodies were cloned into the pET-Sumo vector [22]. The recombinant scFv contains carboxy-terminal HIS-tag and HA-tag used for purification and detection. Antibodies were produced in *Shuffle pLys Y* cells, grown at 37°C, 200 rpm, until an OD (600nm) was achieved. Induction was performed adding 1 mM IPTG (Sigma-Aldrich, cat: I6758). Additional incubation at 200 rpm, 25°C for 4 hours was carried out. Cells were sonicated, and the clarified supernatant was purified by immobilised metal affinity chromatography on 1 ml HISTRAP HP (Cytiva, cat: 17524701) column in AKTA system (Cytiva). The eluted fractions were cleaned and concentrated in Amicon Ultra-2 30 kDa (Merck Millipore, cat: UFC203024) by diluting them in PBS (150 mM NaCl, 10 mM NaHPO4, pH 7.4), and further quantified by Bradford Assay as described in Kielkopf [23]. Antibody expression and purification were evaluated using polyacrylamide gel electrophoresis stained with Coomassie Brilliant Blue G-250 and Western-blot using a mouse anti-HA (Santa Cruz Biotechnology, cat: sc-7392) at a dilution of 1:1,000. Bovine anti-mouse IgG conjugated to alkaline phosphatase (Santa Cruz Biotechnology, cat: sc-2373) was used as the secondary antibody, at a dilution of 1:2,000.

### Binding of Recombinant Antibodies to Biotinylated Viral Peptide

One hundred µL of 240 µM recombinant antibody solution in PBS was adsorbed onto 96-well NUNC MaxiSorp ELISA plate at 4°C overnight. The wells were washed 3 times with 200 µL of PBST (PBS with 0.1% Tween 20) and blocked with 150 µL of 1% BSA in PBST for 1 hour at 37°C. The wells were washed as before, and serial dilutions (3.9 µM; 1.3 µM; 0.43 µM; 0.14 µM in 100 µL of PBS) of biotinylated viral peptide were added and incubated for 1 hour at 37°C. The plate was washed, and 100 µL of alkaline phosphatase-conjugated streptavidin (SeraCare, cat: 475-3000), diluted 1:1000 in TBST, were added and the plate incubated at 37° for 1 hour. After washing the plate with PBST, the captured viral biotinylated peptides were detected with 1 mg/ml pNPP (*p*-nitro-phenyl-phosphate, Thermo Fisher Scientific, cat: 34045) diluted in Diethanolamine substrate buffer 1X (Thermo Fisher Scientific, cat: 34064). The plate was incubated with the pNPP solution at room temperature for 30 min, and the reading performed with a 405-nanometer filter in a SpectraMax M2e spectrophotometer (Molecular Devices). Mean absorbances were determined from replicates of at least two independent experiments, with antibodies from different batch and purification experiments. A mock scFv (anti-DNA) produced and purified under the same conditions of the anti-Zika scFv was used as a negative control.

### Binding to Flavivirus

Approximately 1.5 × 10^2^ PFU/mL (plaque-forming units per millilitre) of inactivated ZIKV and YFV in 100 µL of PBS (pH 7.4) were adsorbed onto a NUNC MaxiSorp ELISA plate at 4°C overnight. The wells were washed 3 times with 200 µL of PBST and blocked with 150 µL of 1% BSA in PBST, for 1 hour at 37°C. ZIKV and YFV sensitised and blocked plates pre-sensitised plates with a mixture of DENV type 1-4, highly purified and in equivalent amounts (Diagnostic kit EUROIMMUN, cat: EI 266a-9601-1 M, pre-coated plates), were used to analyse antibody binding to whole Flavivirus particles. Ten 3-fold serial dilutions (5.93 µM to 2.71 ηM in 100 µL of PBST) of recombinant antibodies were incubated on the virus coated plates for 1.5 hours at 37°C. The plates were washed with PBST, and 100 µL of alkaline phosphatase-conjugated anti-HIS tag antibody (Sigma-Aldrich, cat: A5588), diluted 1:2,000 in TBST, was added and incubated for 1 hour at 37°C. After washing the plate, bound scFvs were detected with pNPP 1 mg/mL, and read at 405 nm in SpectraMax M2e spectrophotometer (Molecular Devices). Mean absorbances were determined from replicates of at least three independent experiments, with antibodies from different expression and purification experiments. The half effective concentration (EC_50_) of antibodies needed for half the maximal binding was determined by non-linear regression analysis in GraphPad Prism. A mock scFv (anti-DNA) was used as a negative control.

### Virus Production and Titration

A culture flask containing 6.0 × 10^6^ *Vero* cells was infected for one hour at 37°C with 0.01 MOI ZIKV in 5 mL of MEM medium with 2% FBS (Serum Fetal bovine; Thermo Fisher Scientific, cat: 12657029). After one hour of infection, 5 mL of MEM medium with 10% FBS and 1X Antibiotic-Antimycotic (Thermo Fisher Scientific, cat: 15240062) was added directly, and the culture incubated at 37°C and 5% CO_2_ until a cytopathic effect was observed, usually for 40 to 48 hours. After this period, the culture was centrifuged at 1,300g at 4°C, and the supernatant containing the virus stored at −80°C.

For ZIKV titration by plaque assay, 1.5 × 10^5^ *Vero* cells per well (24-well plate) were cultured for 24 hours at 37°C and 5% CO2 in 500 µL/well of MEM medium with 10% FCS and 1X Antibiotic-Antimycotic solution. Cells were infected with 100 µL of 10-fold dilutions of virus (10^−1^ to 10^−7^) for one hour at 37°C. The wells were gently covered with 500 µL of MEM semi-solid medium containing 1.5% CMC (Carboxymethylcellulose Medium viscosity; Sigma Aldrich, cat: C4888), 2% SFB, 1X Antibiotic-Antimycotic, and 1% non-essential MEM amino acids (Thermo Fisher Scientific, cat: 11140068). The culture plates were incubated at 37°C and 5% CO_2_ for four days. After this, the semi-solid medium was removed, and the cells were fixed with 500 µL of 10% Formaldehyde overnight. Afterwards, the wells were washed with 1 mL of water, by incubation at 37 °C for 15 min, and then stained with 500 µL of dye solution, Crystal violet 1% (Sigma-Aldrich, cat: C0775), for 30 min at room temperature. Lysis plaque counts and virus concentrations were determined in PFU/mL (plaque-forming units per ml).

### Plaque Reduction Neutralization Assay

The neutralizing activity of the recombinant scFvs was evaluated by the test of plaque reduction neutralization (PRNT) in *Vero* cells. For that, 1.5 × 10^5^ cells per well were cultured for 24 hours at 37°C and 5% CO_2_ in a 24-well culture plate in 500 µL of MEM medium containing 10% FBS and 1X Antibiotic-Antimycotic. Eleven 4-fold dilutions of the recombinant antibodies (12.2 µM to 0.01 ηM), in a volume of 100 µL of MEM medium with 2% FBS, were incubated with 100 µL of 1.24 × 10^3^ PFU/mL ZIKV (tittered for yielding 35 to 45 plates per well in a 24-well plate) for 1 hour at 37°C. The antibody and virus mixtures were added to the cells and incubated for 1 hour at 37°C. The wells were gently covered with semi-solid medium as described above. The culture plates were incubated at 37°C and 5% CO_2_. After four days, the semi-solid medium was removed, the cells fixed and stained as described above, and the plaques counted. The value of half the maximum inhibitory concentration (IC_50_) corresponds to the concentration of recombinant antibody that resulted in a reduction of half the number of plaques observed in the wells where cells were infected without the presence of antibody. IC_50_ was determined by four-parameter non-linear regression in GraphPad Prism. Neutralization values were obtained from the mean of replicates of at least two independent experiments. An anti-DNA scFv produced and purified under the same conditions of the anti-Zika scFv was used as a negative antibody. Controls were prepared with uninfected cells incubated with the highest protein concentration to assess the toxicity of the recombinant molecules.

## Results

### Antigen design and selection

For selecting specific antibodies to ZIKV and other Flavivirus, a peptide corresponding to a conserved region of the FL of Flavivirus was designed to serve as an antigen in selecting a naive Phage Display combinatorial library. The antigen was designed based on the sequence alignment of the E protein domain II of different Flavivirus (**Fig 1A**) and was conceived to contain the FL linked, by a disulfide bridge, to a contacting loop in the domain II region, the SRCPT peptide. The use of the cysteine-bound peptide aims to bring the conformation of the peptide closer to that which occurs in the viral particle (**Fig 1B**).

**Fig 1.**
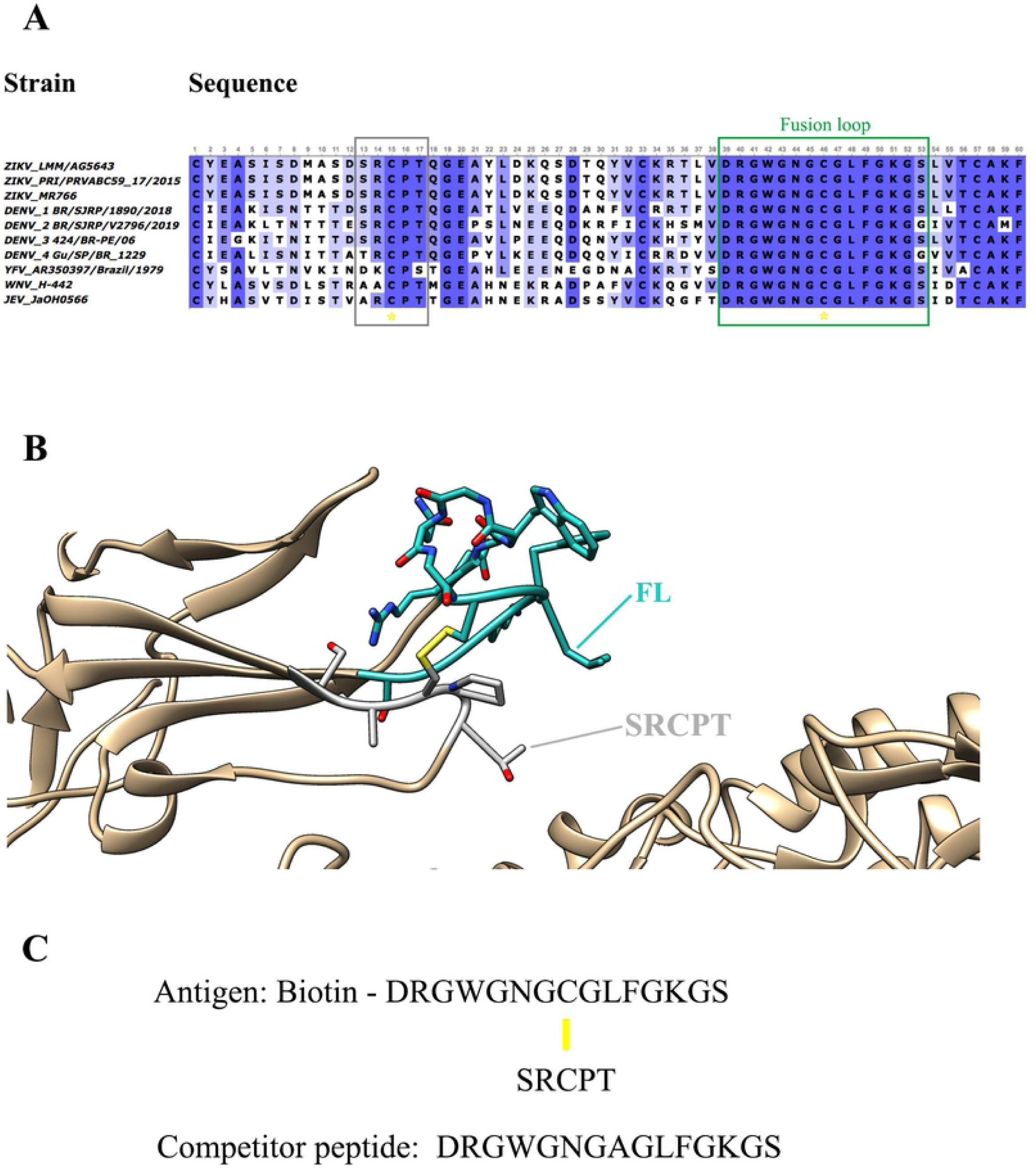
Design of antigen for the selection of antibodies anti-FL. (**A**) Antigen design was based on the alignment of the fusion loop sequences (green rectangle) found in the Protein E Domain II from different Flavivirus strains. A disulfide bridge connects the FL sequence to the SRCPT peptide (gray rectangle). Yellow asterisks highlight the conservation of cysteines. (**B**) Structural representation of the FL (cyan) in the ZIKV protein E domain II. On the viral envelope, FL project outward domain II underpinned by the SRCPT peptide (grey) by a disulfide bridge bond (highlighted in yellow). (**C**) Peptide sequences of the selection antigen with FL, peptide SRCPT and Biotin, and competitor peptide with only FL (with an Ala substituting for Cys) are used for competitive elution. Yellow line: disulfide bridge.

This target peptide was also conjugated to a biotin molecule (biotinylated viral peptide) for binding to the selection plate pre-sensitised with streptavidin. A peptide containing only the FL amino acid sequence (free viral peptide) was also designed (**Fig 1C**) to serve as a competitor in the elution of specific antibodies, in addition to the usual acidic elution (**S1 Fig**). The idea in the competitive elution was to isolate only those antibodies that are bound to FL, avoiding antibodies that bound to the supporting peptide SRCPT, among other unspecific antigens present in the selection platform.

A combinatorial phage-displayed library of human antibody fragments was used for selection against the biotinylated viral peptide. Antigen-specific phages were eluted by an acid solution or by a competitor peptide. Acid and competitive selection were conducted independently throughout the experiment, and analysis of selection progress to understand antibodies’ binding and elution behaviour in the system (**S1 Fig**).

### Analysis of the progress of selection and identification of enriched variable domains

The coding region of VH and VL domains of antibody subpopulations from the original library (before selection) and after selection (acid or competitive elution) (**S2 Fig**) were PCR amplified and subjected to high-performance sequencing (Illumina). The sequences of VH and VL in each round were ordered due to the enrichment of unique sequences, expressed as fold-change (FC), compared to the original library.

Sequencing statistics showed a reduction in the number of unique VH and VL sequences from round 0 (before selection) to round 4 in both Phage Display strategies, indicating a successful selection process (**S1 Table**). The most enriched VH and VL, with the higher fold-change values, were identified. The six most enriched VHs in peptide and acid selection presented FC ranging from 483 to 1,308 and 586 to 3,827, respectively (**Table 1**). The top six enriched VLs after competitive and acidic selection showed FC varying from 80 to 2,308 and 201 to 1,021, respectively. The first most enriched VL in the peptide eluted strategy, Lp1, had FC much higher than the other enriched VLs (**Table 1**).

**Table 1.** Characterization of selected heavy chain variable domains (H) and light chain variable domains (L).

|  | Competitive selection |  |  | Acid selection |  |  |
| --- | --- | --- | --- | --- | --- | --- |
|  | Sequence<sup>a</sup> | FC<sup>b</sup> | V Gene<sup>c</sup> | Sequence | FC | V Gene |
| **VH** | Hp1 | 1,318 | IGHV1-2*02 | Hac1 | 3,827 | IGHV4-34*01 |
|  | Hm1 | 1,074 | IGHV1-3*01 | Hm1 | 1,307 | IGHV1-3*01 |
|  | Hm2 | 746 | IGHV3-66*01 | Hm3 | 838 | IGHV3-7*01 |
|  | Hm3 | 735 | IGHV3-7*01 | Hm2 | 667 | IGHV3-66*01 |
|  | Hm4 | 610 | IGHV3-74*01 | Hm6 | 645 | IGHV3-49*04 |
|  | Hm5 | 483 | IGHV3-7*01 | Hm5 | 586 | IGHV3-7*01 |
| **VL** | Lp1 | 2,308 | IGHK1-33*01 | Lac1 | 1,021 | IGHK1-17*03 |
|  | Lp2 | 395 | IGHK3-20*01 | Lac2 | 711 | IGHK2-30*02 |
|  | Lp3 | 158 | IGHK3-20*01 | Lac3 | 679 | IGHK1-17*02 |
|  | Lp4 | 151 | IGHK2-30*02 | Lac4 | 325 | IGHK4-1*01 |
|  | Lp5 | 85 | IGHK1-33*01 | Lac5 | 318 | IGHK1-17*01 |
|  | Lp6 | 80 | IGHK1-33*01 | Lac6 | 201 | IGHK4-1*01 |

<sup>a</sup>The letters "p", "ac" and "m" indicate variable domain from competitive selection, acid selection or both selections, respectively.
<sup>b</sup>FC values correspond to frequency variation of an unique VH peptide along selection procedure.
<sup>c</sup>V gene family of each sequence was identified based on IgBlast annotation.

Both selections observed top enriched heavy chain variable domains, except for the most enriched VH, Hp1, and Hac1, which were selection exclusive. Hac1 shows an FC de 1,318 after acidic elution and Hp1 with FC de 3,827 after competitive elution. Unlike VH, none of the most enriched VL was shared by both selection procedures (**Table 1**).

The two selections shared the majority of the most enriched VH gene families except for IGHV1-8 and IGHV3-21, exclusive to the acidic selection, and IGHV1-2, exclusive to competitive selection (**Fig 2A**). The IGKV1-33 gene family, overrepresented among top enriched VLs in the competitive selection, was not selected after acidic elution. On the other hand, the IGKV4-1 gene family, the most frequent family among the VLs of acidic selection, only appears in this selection (**Fig 2B**).

**Fig 2.**
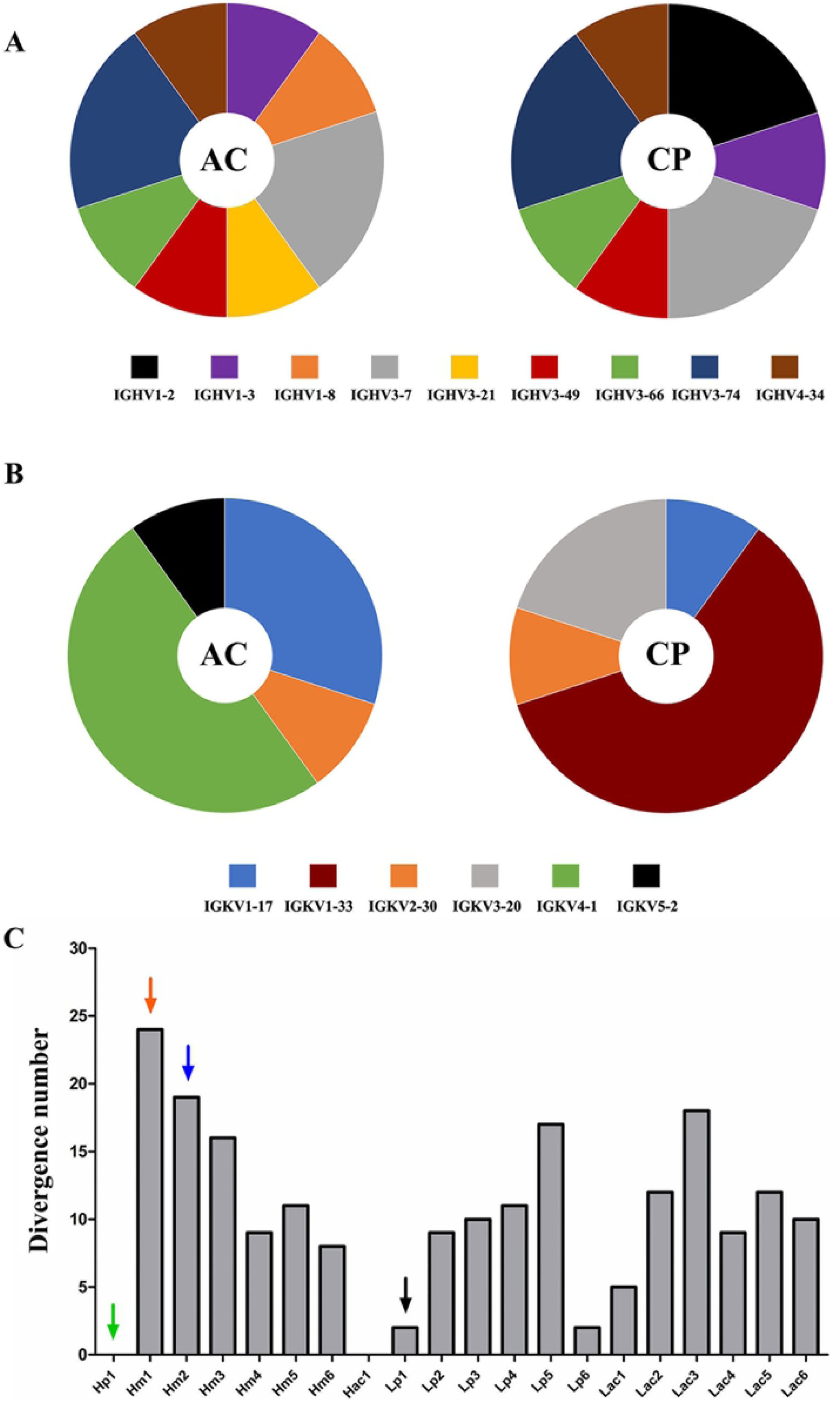
Differences between the most enriched sequences in both selection procedures. Representativeness of the V gene segment families to which the ten most enriched VH (**A**) and VL (**B**) sequences belong, in acid (AC) and competitive selection (CP). The width of each slice of the graph is proportional to the number of sequences belonging to the specific gene family. (**C**) Analysis of the dissimilarity to germline V gene of the most enriched V domains. The degree of dissimilarity of the VH (H) and VL (L) comparing their closest germline sequences is represented by the number of non-identical residues. The arrows mark the three VH and the VL used in the tested scFv.

Antigen-selected VH sequences were more divergent to germline than VL sequences (**Fig 2C and Fig 3**), but variable gene segments in the germline configuration are only found among VH. Hp1 and Hac1, the most enriched VH after each elution strategy, utilise the germline IGHV1-2*02 and IGHV4-34*01 gene segments, respectively. Oppositely, Lp1, the most enriched VL after competitive selection, has two amino acid changes. Other enriched sequences had a high degree of sequence divergences, such as VH Hm1 and Hm2, with 24 and 19 amino acid substitutions compared to the hitherto germline, respectively (**Fig 2C**). This shows enrichment of sequences with both high and low degrees of dissimilarity compared to their germinal counterpart.

**Fig 3.**
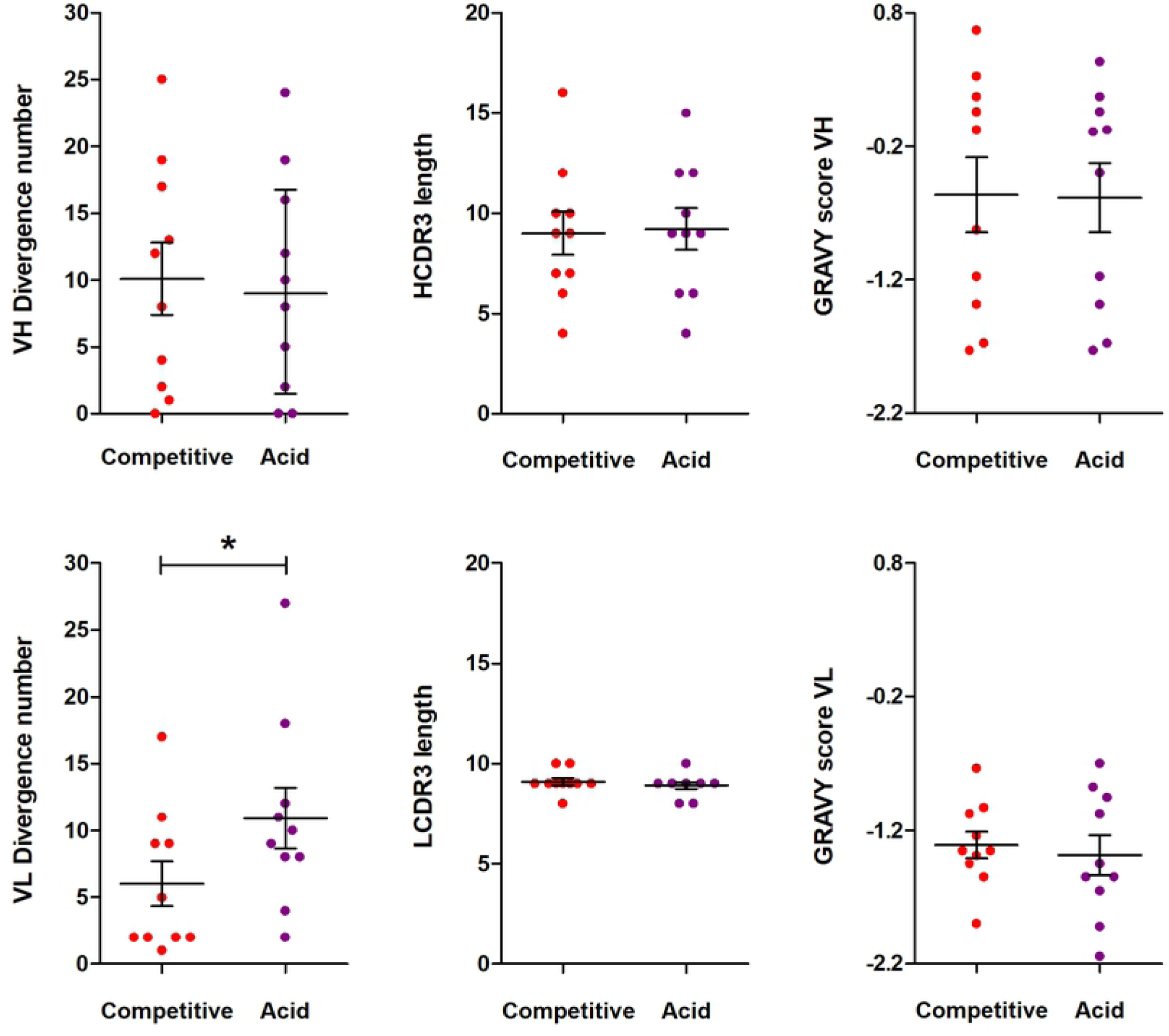
Properties of the most enriched VH and VL. The characteristics of the ten most frequent variable domains in the acid selection (Acid) and competitive selection (CP) were analyzed. The characteristics of VH and VL studied were: divergence to the germline V gene (number of non-identical residues); CDR3 length (number of residues); and GRAVY scores of hydrophobicity of the CDR3 amino acid sequence. Data are represented as the means and SEM in each group (N=10/group). * Means of groups were statistically different at *P < 0*.*05*; unpaired t-test.

The use of either elution strategy leads to VH selection with a varying degree of V gene maturation, CDR length, and hydrophobicity (**Fig 3**). However, for VL the degree of divergence in relation to the germinal sequences is lower in antibodies selected after competitive elution (*p < 0*.*05*).

### Construction of the recombinant anti-Flavivirus antibodies

Top enriched VH and VL domains were strategically combined in scFv (single-chain variable fragment) format, considering their enrichment, presence in both selections or just one (prioritising the competitive selection), and their germinal dissimilarity degree. Three scFv molecules were constructed and analysed: AZ1p, containing Hp1; AZ3m, with Hm1, and AZ6m, with Hm2. The exceptionally enriched Lp1 was chosen as the VL counterpart of all three scFv **(Fig 4)**. Another set of scFv molecules was constructed utilising another VL, Lac1. However, these antibodies showed low expression levels (data not shown) and were not further analysed.

**Fig 4.**
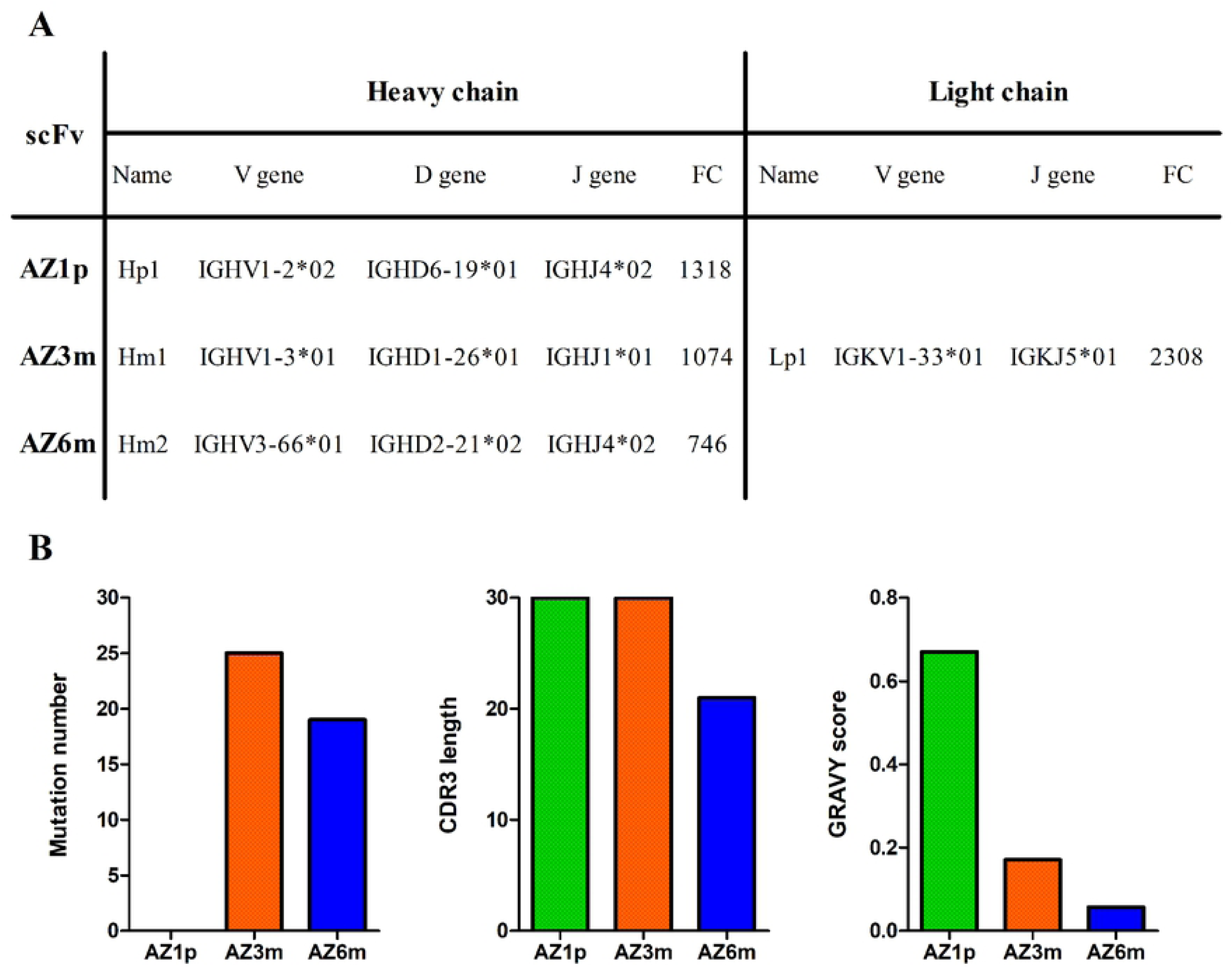
Recombinant scFvs features. Three anti-FL monoclonal antibodies, AZ1p, AZ3m and AZ6m were constructed, by combining the enriched VHs (Hp1, Hm1 and Hm2) with the most enriched VL (Lp1), respectively, in an scFv format (A). Characteristics of the three scFvs, considering the variable domain of the heavy chain (B). Profiles of VH amino acid divergence (compared to their germline sequence) and length and hydrophobicity of the CDR3 protein sequence show differences between tested antibodies.

Recombinant antibodies were expressed and purified for further characterization (**S3 Fig**). The three scFvs generated remained the ability to bind to the synthetic antigen used in the Phage Display selection, showing different binding profiles. AZ6m antibody showed the highest binding activity to the viral antigen, binding at least twice higher than observed with the other scFvs (**Fig 5**).

**Fig 5.**
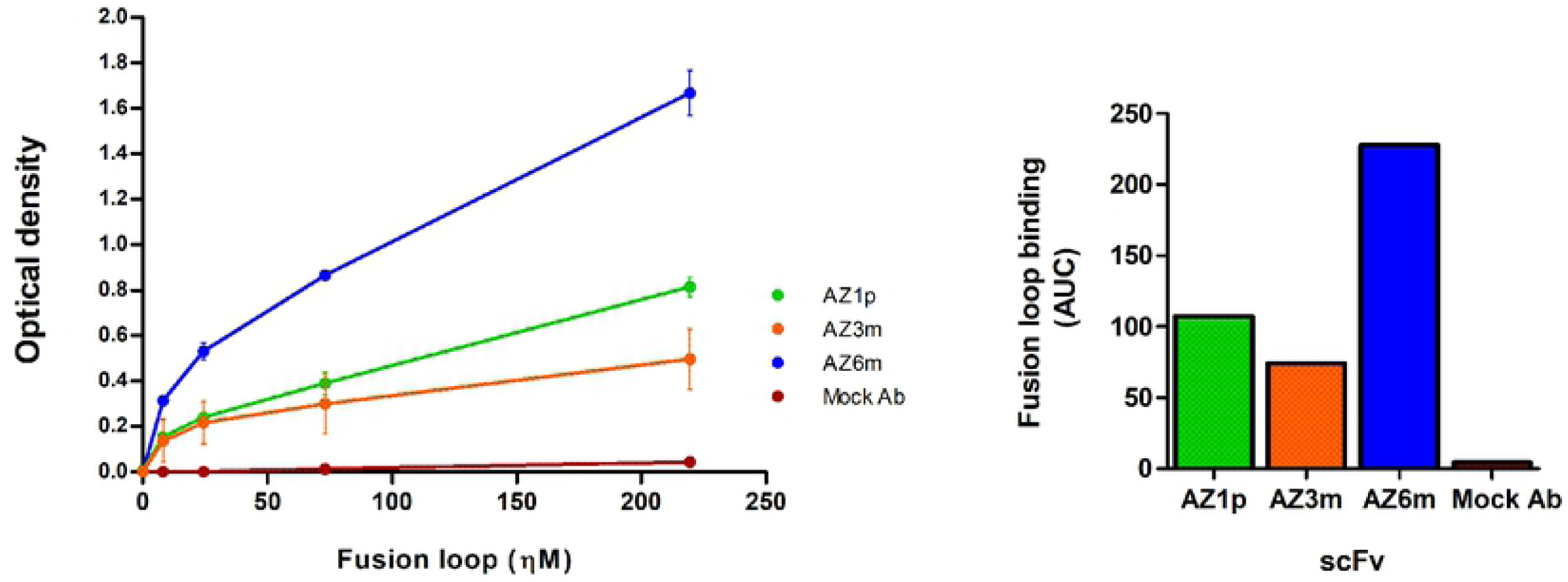
Binding of monoclonal antibodies to the fusion loop antigen. On the left, the binding of the recombinant scFvs to the fusion loop used in the selection was analyzed by immunoassay (ELISA). The normalized area under curve for each antibody FL binding is on the right. Data are presented as mean +/− SD. An unspecific scFv, produced in the same conditions of the anti-FL scFvs, was used as a negative antibody (Mock Ab).

### Binding to different Flavivirus

To determine whether the generated antibodies reacted to FL in their native conformation in different Flavivirus, they were screened for binding to ZIKV, YFV and the 4 DENV serotypes using viral particles. All antibodies showed some binding capacity to all Flavivirus tested, following the same order of binding activity among Flavivirus (**Fig 6A-6C**). Contrasting with peptide binding, AZ1p scFv showed the highest virus binding activity, with EC_50_ values of 14.67; 16.24 and 545.8 ηM, followed by AZ6m, with EC_50_ values of 361.7; 256.4 and 740.0 ηM, and finally AZ3m, with values of 2,479.0; 974.8 and 1,094.0 ηM for ZIKV, YFV and 1-4 DENV respectively (**Fig 6D**).

**Fig 6.**
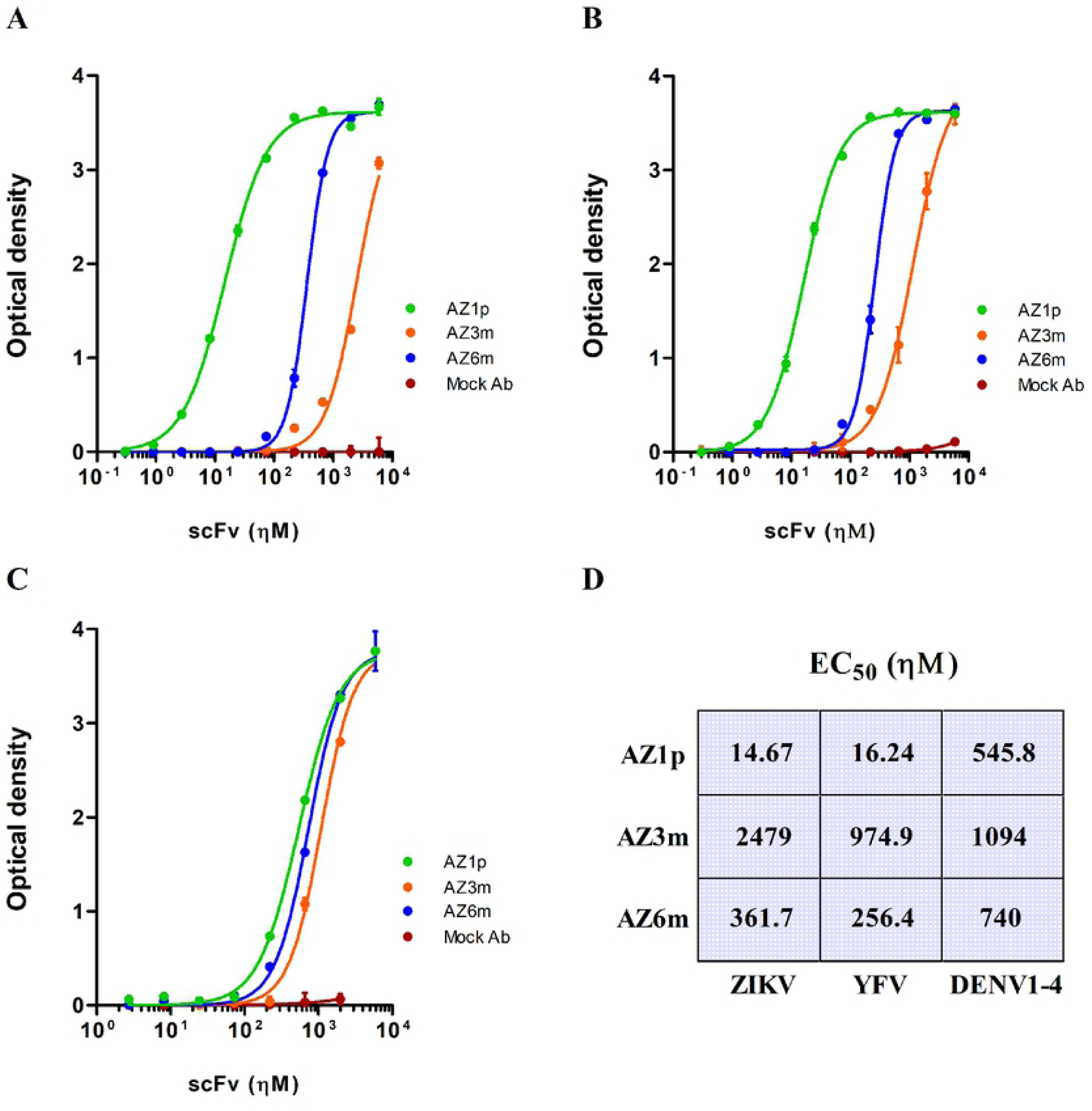
Anti-flavivirus binding activity of the selected scFvs. Recombinant scFvs (AZ1p, AZ3m and AZ6m) were analyzed for their ability to bind to different flavivirus: ZIKV (**A**), YFV (**B**) and an equal mix of the four DENV serotypes (**C**). In these experiments, the plates were sensitized with the whole viral particles. The half effective concentration (EC_50_) for each scFv against each flavivirus were also determined (**D**). Data are presented as mean +/− SD. An unspecific scFv was used as a negative antibody (Mock Ab).

The binding pattern to ZIKV and YFV was similar. The differences between antibody binding to DENV serotypes were minimal and were smaller than that observed for ZIKV and YFV. The highly divergent AZ3m presented a very low binding capacity to ZIKV. However, the antibody AZ1p, with a germinal VH, showed expressive binding to ZIKV and YFV (**Fig 6A and 6B**).

### Neutralizing activity to Zika virus infection

A plaque reduction neutralization test (PRNT) with the Zika Virus was performed to assess whether the constructed anti-FL scFv has neutralizing capacity. The AZ1p and AZ6m monoclonal antibodies showed similar neutralizing activity, with half-maximal inhibitory concentration, IC_50_ of 397.4 ηM and 311.4 ηM, respectively (**Fig 7**). The AZ3m antibody did not show neutralizing capacity (IC_50_ > 100 µM), corroborating the low binding activity to the ZIKV particle.

**Fig 7.**
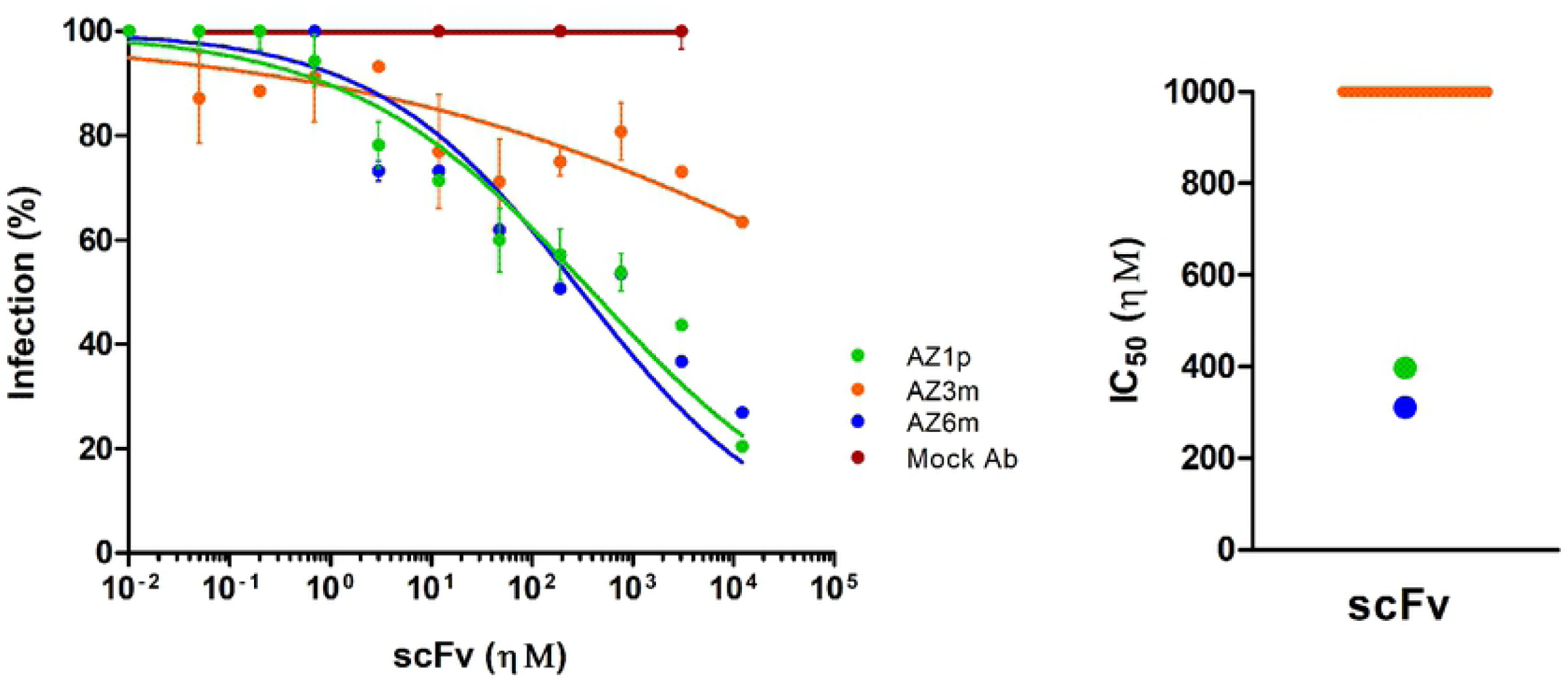
Neutralization of Zika virus infection. The ability of scFvs to neutralize Zika virus was measured in a plaque reduction neutralization test (PRNT). On the left, the percentage of infection was calculated by the ratio between the number of plaques formed in the presence and absence of antibody. On the right, half-maximum inhibitory concentrations (IC_50_) represent means of replicates from at least two independent experiments. The representation of the IC_50_ values was limited to 1 µM. The IC_50_ value of the AZ3m greater than 1 µM was plotted as the limit (orange line). An unspecific scFv was used as a negative antibody (Mock Ab).

## Discussion

Antibodies specific to FL may have important neutralizing potential to infections of different strains of Flavivirus [24]. In this work, we reported the productive selection of anti-FL antibodies, reducing the library diversity throughout the rounds, warranting the selection process. The selection leads to the identification of unrelated VH and VL enriched in response to the selection. Interestingly, both elution schemes lead to a common set of VH among the top selected, except for the first most enriched VH: Hp1 and Hac1. Unlike heavy chain domains, none of the most enriched VLs were selected in both schemes. The most significant differences between the selections are found in the VLs: in the FC values between the enriched VLs, the V gene families, and the degree of dissimilarity to the germinal sequences. This corroborates the idea that distinct elution schemes lead to the selection of particular antibodies and also shows that different pressure determinants drive the selection of VHs and VLs. Thus, the elution protocol impacts, at least in part, the observed selected antibodies.

Some of the selected VH and VL showed a high dissimilarity with their germline sequences. Likewise, studies of the antibody-mediated immune response to Influenza virus and HIV have shown that it is possible to accumulate a vastly hypermutated VH in highly neutralizing antibodies [25,26], which may explain the nature of these highly hypermutated sequences. However, a naive phage library is not supposed to contain antigen-driven hypermutated antibodies, and it is considered a limitation for using such libraries. Even though the library used in this work may be regarded as naive for viral infection, the library used here was assembled from donors from an endemic area for DENV. Considering that prior contact events with other Flaviviruses can lead to protective immunity against ZIKV infection [27,28], preexisting highly hypermutated VH can be selected that bind to conserved Flavivirus antigens, such as observed in AZ6m.

Along with hypermutated VH, in both selections, the highest enrichment went to germline VHs (Hp1 and Hac1). Germline sequences are generally associated with low-affinity antibodies, but some V genes seem to possess an intrinsic antigen recognition [29,30]. Moreover, it has been reported that germline-like antibodies from convalescent individuals have a potent neutralizing capacity against ZIKV and SARS-CoV-2 [31,32]. Indeed, the scFv AZ1p, which contains Hp1, efficiently neutralized ZIKV. Thus, we showed that both germinal-like antibodies, such as AZ1p, and antibodies with a high number of variations, such as AZ6m, may have neutralizing potential for viral infections.

The therapeutic antibody design influences its pharmacological properties. ScFv lacks Fc, which makes ADE reaction improbable, and it has a shorter half-life and smaller molecular size. These aspects lead to a greater ability to penetrate biological tissues, such as neural tissues, and antigenic structures, as cryptic antigens such as FL [33,34]. On the other hand, a bivalent antibody may have more significant binding and neutralization capabilities. Sharma and colleagues showed that a highly cross-reactive antibody, C10, increases its binding and neutralization to ZIKV and the four types of DENV when it changes from a Fab format to a complete IgG [35].

A major challenge in developing antibodies selected with mimetic peptides is binding to the antigen in its native conformation. Moreover, the FL is a conserved viral epitope but also not easily accessible. Therefore, anti-FL antibodies have been thought to have a lower neutralizing capacity [36]. Thus, the antibodies were tested for binding to Flavivirus, ZIKV, YFV, and DENV, using whole viral particles to assess these hypotheses. All scFv antibodies were able to bind to the structurally conserved FL of all tested viruses. However, binding to DENV may be underestimated. The four serotypes of DENV were tested together, which could hinder the clear differentiation of the antibodies’ binding ability to each serotype. Despite this, binding to all serotypes is expected due to the high conservation of the FL and the structural similarities between DENVs [37].

Recently, a humanised anti-FL antibody, ZAb_FLEP, showed significant IC_50_ values in PRNT against different strains of ZIKV and DENV [38] exemplifying the high cross-neutralization capacity of monoclonal antibodies. The neutralizing ability of some anti-FL antibodies can be explained by the dynamism of the “viral breathing” process associated with the conformational changes induced by the antibody binding leading to the exposure of FL, even in a context without acidification [39,40]. The neutralizing potentials observed from AZ1p (397.4 ηM) and AZ6m (311.4 ηM) were better than other previously described anti-fusion loop antibodies as 2A10G6 with IC_50_ = 1,67 µM [41,42] and C5, IC_50_ > 10 µM [43], and scFv that binds to antigens outside FL, such as scFv45-3, anti-DIII, IC_50_ = 461 ηM [44] and m301, anti-DIII, IC_50_ ∼2 µM [45]. Therefore, this work showed that anti-FL antibodies can be neutralizing. Even though we did not test AZ1p and AZ6p for neutralization of other Flavivirus, their significant EC_50_ for YFV and DENV suggest them as *bonafide* pan-Flavivirus neutralizing antibodies.

The development of new neutralizing antibodies and the combination of antibodies with different specificities are needed to increase treatment efficiency of infections, avoiding the persistence of a high viral load and the possibility of virus immune scape by mutation [46]. The successful combination of different antibodies has already been demonstrated for infection with Zika virus [46], with Ebola virus [47] and with HIV [48]. Since immune evasion by viral mutation were observed in non-human primates treated with anti-ZIKV antibodies with potent binding and neutralizing capacities [49,50], it is essential to dispose of a panel of antibodies targeting key conserved regions of the envelope, the main contribution of this wok.

Data here report two novel neutralizing antibodies AZ1p and AZ6m, representing an important perspective for the treatment of ZIKV infection and potential interventions in infections by other Flavivirus. They can be used as a neutralizing strategy against different strains and variants using a single drug, overcoming the immunological escape and possible ADE reaction. Future *in vivo* neutralization assays with ZIKV, YFV and DENV may attest to the therapeutic potential of these antibodies.

## Acknowledgments

We are grateful to Prof. Connie McManus for English revision.

## Supporting information

**S1 Fig. Schematic representation of the selection processes**. A combinatorial library was used to generate fusion phages expressing human antibody fragments on their surfaces. Four selection rounds were performed, increasing the stringency at each round by raising the number of washes. Two strategies eluted the specific phages. One used an acidic solution to disfavour antibody-antigen binding. The other used a competitively saturating solution of the free unlabeled FL antigenic peptide. Eluted phages were amplified separately for a new selection round.

**S2 Fig. Generation of variable domain genes amplicons**. Phagemid pools from the original library and the fourth round of selection were used as templates for PCR using specific primers for VH (capital letters) and VL (lowercase letters) domains amplification. The amplicons were obtained through antibody library before selection (A, a) and from the fourth round of selection using acid (B, b) or competitive (C, c) elution. M: 100 bp DNA Ladder.

**S3 Fig. Analysis of IMAC purified recombinant scFvs**. The scFv were expressed in bacteria and purified by affinity chromatography in nickel columns. The production of antibodies AZ1p, AZ3m and AZ6m were analysed by denaturing SDS-PAGE (**A**) and detected with anti-HA tag antibody by western-blot (**B**).

